# MRI-free OPM-MEG recovers medial temporal lobe theta during scene imagination

**DOI:** 10.64898/2026.08.10.743897

**Authors:** Conor Thornberry, Prathiksha Math, Margarida Cohen Serra, Robert Seymour, Clodagh Nolan, Robert Whelan

**Affiliations:** School of Psychology, Trinity College Dublin, Dublin, Ireland; Trinity College Institute of Neuroscience, Trinity College Dublin, Dublin, Ireland; Department of Physiology, School of Medicine, Trinity College Dublin, Dublin, Ireland; University of Oxford, Oxford Centre for Human Brain Activity.

**Keywords:** Optically pumped magnetometers, medial temporal lobe, hippocampus, theta, scene imagination, template warping

## Abstract

Optically pumped magnetometer magnetoencephalography (OPM-MEG) offers a wearable, movement-tolerant alternative to conventional cryogenic MEG, placing sensors closer to the scalp and, in principle, improving sensitivity to deep sources. This is advantageous for examining subcortical structures that are affected by ageing, disorders and disease, such as the hippocampus. However, it remains unclear whether previously reported activity (such as the attenuation of theta oscillations during the imagination of novel scenes) can be recovered from the medial temporal lobe (MTL) with OPM-MEG, and whether an individual structural MRI is required for accurate source localisation. Here, fifteen adults completed a scene imagination task. Initially we applied a 12-parameter template warping coregistration pipeline to the full sample. Following source reconstruction, we recovered the expected attenuation of theta (4-8 Hz) power during scene imagination compared to a counting baseline, with a significant cluster of activity peaking in the left parahippocampal gyrus. The cluster’s centre of mass was localised to the left hippocampus (*t* = -3.55, *p* = 0.048, whole-brain FWE-corrected) and was mostly confined to the left medial temporal lobe. We further supported our findings by using an individual T1-weighted MRI reconstruction pipeline in six participants who had these scans available. The two approaches produced similar whole-brain topographies and localised the peak MTL theta effect to left hippocampus, with temporal-lobe conjunction centroids 3-mm apart. These findings provide evidence that the theta attenuation of the scene construction network can be recovered at the group level with OPM-MEG, without an individual MRI.

## 1. Introduction

Optically pumped magnetometers (OPMs) can measure femtotesla scale changes in magnetic fields and have the potential to become the leading technology for magnetoencephalography (MEG) research in neuroscience (Brookes et al., 2022). Conventional MEG relies on Superconducting Quantum Interference Devices (SQUID) sensors, which require liquid helium to maintain superconductivity. This cryogenic cooling makes systems large, expensive and less flexible, as the equipment includes a rigid helmet containing the sensors. As the helmet cannot adapt to individual head sizes, the gap between the sensors and the scalp reduces signal strength, compromising accuracy. Additionally, the static nature of sensor placement restricts mobility, limiting the study of naturalistic behaviours (Brookes et al., 2022). Optically Pumped Magnetometer MEG (OPM-MEG) addresses these constraints (Schofield et al., 2022). OPM-MEG provides a wearable flexible system mounted on a 3D-printed cast placed over the participant’s head, positioning the sensors closer to the scalp (Brookes et al., 2022). This, combined with field nulling (Holmes et al., 2018), allows for natural movement during recording, making OPM-MEG particularly suitable for studying ecologically valid behaviour in humans (Boto et al., 2018; Zabbah et al., 2026). In addition, the increased proximity of the sensors improves sensitivity to neural signals. This in turn, enhances the ability to investigate activity in deeper brain regions, such as the hippocampus, which is central to several cognitive processes (Maguire et al., 2016). For example, spatial navigation relies on naturalistic movements, deep brain structures (such as the hippocampus) as well as oscillatory brain activity thought to be generated from these deep sources (Buzsáki & Moser, 2013; Chersi & Burgess, 2015; Ekstrom & Hill, 2023).

Reconstruction of medial temporal lobe and hippocampal sources has been achieved using conventional MEG over the last number of years. This involves reducing the distance between the head and the sensors, as well as the utilisation of hippocampal-dependent tasks (Clark et al., 2018; Hassabis et al., 2007). Using conventional MEG, Barry et al. (2019) successfully reported left hippocampal theta power modulation during novel scene construction alongside the ventromedial prefrontal cortex (vmPFC). Scene construction is thought to be carried out automatically by the hippocampus, generating a contextually relevant spatial representation that may serve as the early foundation for the formation of episodic memory (Bertossi et al., 2016; Maguire & Mullally, 2013). Decreases in temporal lobe theta power have been demonstrated in MEG, scalp and intracranial EEG during episodic memory recall (Greenberg et al., 2015; Herweg et al., 2020; Long et al., 2014), spatial learning (Thornberry et al., 2023) and scene discrimination (Lucie-Read et al., 2024). Herweg et al. (2020) suggest that decreases in theta could be mechanistically related to increases in task-engagement, which would support the idea of hippocampal scene construction acting as contextual framework from which learning and memory is built (Zeidman & Maguire, 2016). Imagination therefore captures the formation of this initial spatial schematic via the hippocampus and vmPFC (Monk et al., 2021). The consistently observed neural signal of theta attenuation in the hippocampus makes it a valuable target to assess whether OPM-MEG can recover Medial Temporal Lobe (MTL) activity, as well as the extended neural system proposed to be involved (Aggleton & O’Mara, 2022; Lucie-Read et al., 2024).

To date, only one OPM-MEG study has attempted to examine this effect. Barry and colleagues (Barry, Tierney, et al., 2019) successfully source localised right hippocampal theta attenuation in three individuals during a novel scene imagination task using OPM-MEG. They also recovered comparable, left-lateralised theta attenuation with conventional MEG in the same individuals. These data were consistent with previous findings from fMRI (Dalton et al., 2018; Hassabis et al., 2007) and SQUID MEG work (Barry, Barnes, et al., 2019; Clark et al., 2018; Zeidman & Maguire, 2016). However, Barry et al. used 24 single-axis OPM sensors which was the state of the art at the time, but the technology has developed since. Correct positioning of OPM sensors is vital for the recovery of hippocampal signals, as well as accurate source reconstruction and minimal coregistration error (Zetter et al., 2018). These limitations, alongside a small sample, typically would require clean data (Seymour et al., 2022), very accurate forward models (Henson et al., 2009) and heavily rely on the use of an individual T1 MRI scan (Gohel & Khare, 2023).

Most recently, it has been shown that cortical activations measured using OPM-MEG are recoverable without an individual MRI. Rhodes et al. (2025) successfully reconstructed induced beta band responses in twenty individuals with relative accuracy using a head-matched template MRI. However, it is still unclear whether previously reported task-related signals can be investigated at the group level from deeper sources, such as the hippocampus. Reducing the need for structural MRI scans, even for subcortical group-level investigations, is highly advantageous for studying cognitive processes in groups for whom an MRI may not be tolerated, such as children or patients. In addition, hippocampal atrophy is an established marker of Alzheimer’s disease (Apostolova et al., 2006; Aumont et al., 2025). Without MRI scans, this structure and its contributions can be studied in patient cohorts during naturalistic, movement tolerant OPM scanning.

Therefore, we set out to investigate whether MRI-free, template-based OPM-MEG could recover scene construction theta power attenuation (4-8 Hz) from the human MTL. Fifteen adults completed a scene-imagination task. In six subjects who had a structural scan, we compared template-based reconstruction against individual-MRI reconstruction to verify that the template pipeline results were sensible. Relative to Barry et al., we leveraged a triaxial 60-sensor array, with improved temporal sensor placement. This resulted in taking one sensor from the frontal (AFz) and one from the back (P7) to fill slots at the temporal regions which for our setup, are typically left empty. Finally, we also used an increased number of trials, all of which should reduce error and yield clearer group-level signal from subcortical sources (Pu et al., 2018; Zetter et al., 2018). We hypothesised that we would find a significant theta power decrease during scene imagination relative to a counting baseline localising to the medial temporal lobe. Given the right lateralisation from Barry et al.’s original OPM study, we hypothesised a right-hippocampal peak (Barry, Tierney, et al., 2019).

## 2. Methods

### 2.1 Participants

Fifteen participants completed the OPM-MEG experiment (11 female; mean age 26.9 ± 2.6 years, range 21.9 - 30.3 years). Of these, six also underwent a T1-weighted structural MRI (3 female; mean age 27.5 ± 2.3 years). The remaining nine completed the identical OPM-MEG protocol without an MRI. All participants had normal or corrected-to-normal vision and no history of psychiatric or neurological disorders. OPM-MEG data collection took place at Trinity College Dublin. The research protocol was approved by the Trinity College School of Psychology Research Ethics Committee (Reference: 5803) and written informed consent was obtained from all participants.

### 2.2 Experimental Paradigm

Participants performed a scene imagination task during OPM-MEG scans. This involved the imagination of novel scenes in response to single scene words presented one at a time (e.g. “casino”, “boardroom”). The scene stimuli have been used in previous fMRI (Clark et al., 2018), SQUID-MEG (Barry, Barnes, et al., 2019) and OPM-MEG (Barry, Tierney, et al., 2019) experiments and were rated as both highly imageable and scene-evoking by at least 70% of an independent sample of participants (Clark et al., 2018). Participants were also presented with a baseline condition, during which number stimuli were presented, in English on the screen (e.g., Eighteen; Twenty-seven). These number stimuli were matched to the scene words by the quantity of letters and syllables.

During scanning, experimental stimuli were projected onto a screen inside the magnetically shielded room using a Propixx projector (Vpixx Technologies, Inc: Saint-Bruno, QC) running at a 480 Hz refresh rate and a 960 x 540 resolution. The screen measured 51.5 x 29.5 cm and was positioned at a viewing distance of ∼55 cm. To prepare participants for each trial type, they were first cued with either the word “scene” or “counting” (Figure 1). Participants were then presented with the word for 2000 ms and instructed to close their eyes on the auditory cue which was presented following a delay of 100ms. During scene trials, participants constructed a novel, vivid scene in their imagination based on the cue (e.g. “casino”). Counting trials involved mentally counting backwards in threes from a number cue (e.g. “ninety-one”). The task periods were 3000 ms in duration. Participants then heard a second beep, opened their eyes and were presented with a question displayed on the screen in front of them to which they responded using a MEG-compatible response box. They were asked to rate whether or not they were successful in the scene imagination task, and whether or not they paid attention during the counting trials. Following this, there was a 1000-ms delay before the next trial. Only trials in which participants confirmed that they were successful were subsequently analysed.

**Figure 1:**
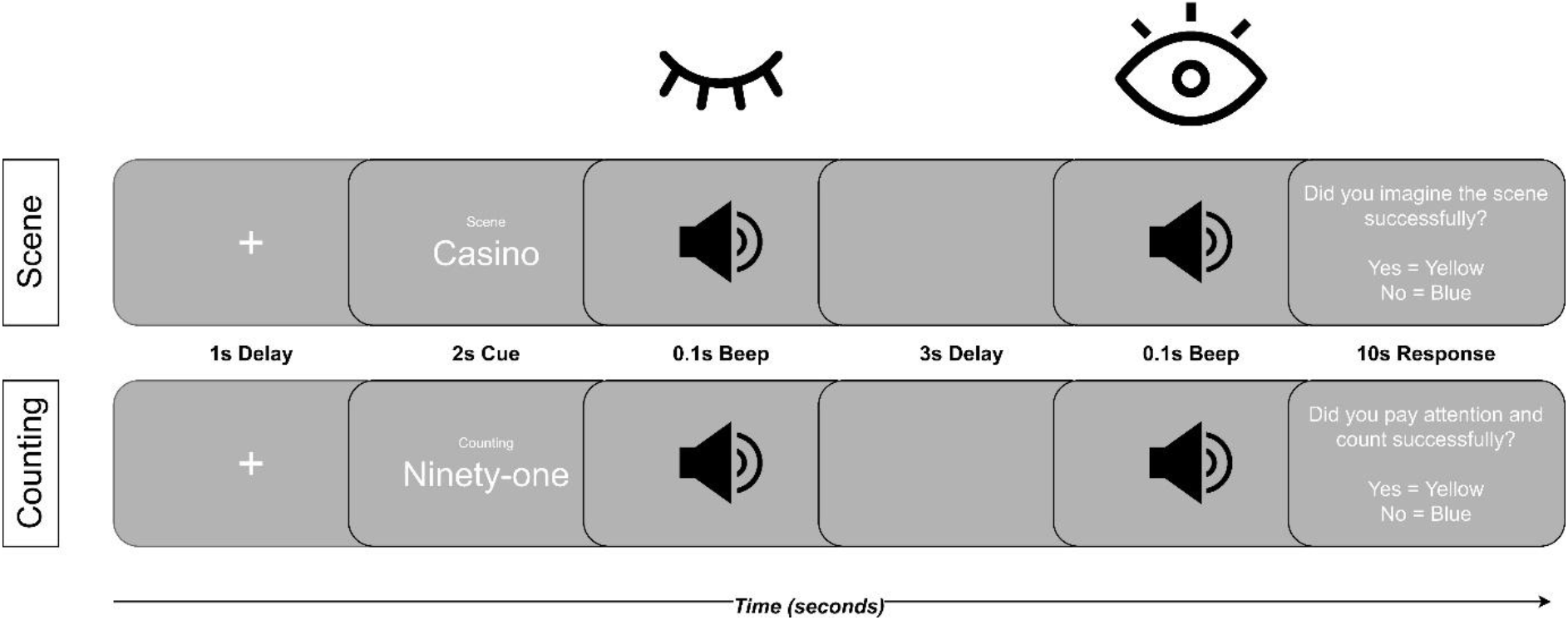
Experimental design for both task conditions. Schematic of a single trial for each condition. Each trial began with a visual cue indicating the upcoming condition (“scene” or “counting”), followed by a stimulus word presented for 2000 ms: a scene-evoking noun (e.g. “casino”) on scene trials or a number word (e.g. “seventeen”) on counting trials. After a 100 ms delay, an auditory cue prompted participants to close their eyes and perform the task for 3000 ms: constructing a novel, vivid mental scene (scene trials) or mentally counting in threes from the cued number (counting trials). A second auditory beep signalled participants to open their eyes and respond via an MEG-compatible response box, rating whether they had successfully imagined the scene or attended to the counting. A 1000-ms inter-trial interval followed. Number stimuli were matched to scene words for letter and syllable count.

This paradigm is also described in Barry et al. (2019) and most recently in Zabbah et al. (2026). In our version of this task, word cues were delivered visually on-screen following the appearance of a fixation cross. Stimulus presentation was split across two blocks of 100 trials each, with blocks approximately 15 mins in length, with participants given a short break between blocks. Scene and counting trials were drawn from two fixed word lists presented in counterbalanced order across participants, with a short rest at the midpoint. Within each list, trial order (and therefore condition order) was fully randomised. Complete data were not obtained for all participants, as for a small number of cases the OPM recording terminated before task completion. This was due to extreme field-nulling drift in one case. In other cases, data files overwrote earlier recorded signals during the same session. Because starting blocks were counterbalanced, trials lost to incomplete sessions were distributed across both word sets. All successful trials were retained, with scene and counting trial counts equalised within participants by dropping excess trials using MNE’s minimum-timing method, which retains the trials that minimise timing differences between conditions.

### 2.3 OPM-MEG Data Collection

MEG data were acquired using a Cerca OPM-MEG system (Cerca Magnetics, Nottingham, UK), which uses OPMs to measure the femtotesla-scale magnetic fields originating from neural activity (Schofield et al., 2025). To ensure the high-stability environment required for these sensitive quantum magnetometers, all recordings were conducted within a dedicated MuRoom® (2.7 x 2.7 x 2.2 m^3^), which limits any external magnetic fields that would saturate or add noise to the sensor output (Hill et al., 2020; Schofield et al., 2025). This magnetically shielded room (MSR) provides essential multi-layered passive shielding. Furthermore, the walls of the MSR integrate degaussing coils, which provide active shielding and are responsible for demagnetising the inner MuMetal layer, thereby removing remnant magnetisation and reducing the internal residual field (Hill et al., 2020). Additionally, a cCoil control system, consisting of an array of 83 electromagnetic field-nulling coils, provides active magnetic field compensation by generating equal and opposite cancellation fields and gradients. These systems are critical for reducing the spatial variation of the residual magnetic field, ensuring that triaxial sensors remain within their narrow operational dynamic range and maintaining the stability of the zero-field environment (Boto et al., 2018).

Neural signals were recorded using an array of 60 triaxial QuSpin QZFM OPM sensors, which require a near-to-zero background field. Each sensor head is triaxial, measuring the magnetic field simultaneously in three orientations: X and Y (tangential) and Z (radial). The sensors contain a fragile head with a laser; a glass cell filled with rubidium-87 vapour and other optical components. During operation, the internal cell is heated to 150°C and insulated by an aerogel layer to ensure the outer surface temperature remains below 41°C for participant safety. Each sensor head is connected via a flexible ribbon cable to a dedicated electronics module, which is connected to both the Data Acquisition unit (National Instruments cDAQ 9179) and the Acquisition PC. A 3D-printed helmet, corresponding to an average adult size, was used in all experiments and woollen inserts were used to provide additional insulation between the sensors and the participant’s head. Finally, two additional triaxial OPMs were used as a reference array, placed on either side of the participant’s head to continuously monitor the remnant background field and provide feedback for active shielding throughout the session.

### 2.4 Background Magnetic Field Control

The experimental protocol for the OPM-MEG recordings followed a rigorous sequence of hardware initialisation, environmental optimisation, and sensor calibration (Boto et al., 2022; Brookes et al., 2022; Schofield et al., 2024). Once the acquisition and stimulus PCs were operational, the MuRoom® door was closed to perform a 60-second degaussing sequence, which used an AETECHRON amplifier. Following degaussing, static field nulling was performed using the cCoil system and the reference OPM array, and an output gain of 0.33x (0.9 V/nT) was applied to the reference sensors to measure the remnant background field without saturation. Once the background field was stabilised, dynamic field nulling was performed to provide continuous active compensation for real-time field changes throughout the duration of the scan. With the environment stabilised, the OPMs in the helmet array were initialised via the QuSpin UI. Neural data acquisition was recorded via software at a sample rate of 600 Hz with the sensors set to an OPM gain of 1.00x (2.7 V/nT).

### 2.5 Preprocessing & Interference Suppression

For preprocessing the OPM data, we closely followed the OPM-FLUX pipeline (Rakshit et al., 2026) and used MNE-python (Gramfort et al., 2013), the OSL-ephys toolbox (van Es et al., 2025) and FSL (Jenkinson et al., 2012). Data were recorded at 600 Hz and downsampled to 300 Hz. Data were plotted for visual inspection. Flat or excessively noisy sensors were identified by visually inspecting the power spectral density (PSD) across the sensors and examining single sensors that were above the median value of the PSD. These were subsequently marked as bad and removed. Automated artifact detection was performed on sensors using *bad_segments* from *osl_ephys*. The continuous data were evaluated in 300-sample windows using a cross-channel kurtosis metric. Windows with outlying kurtosis values exceeding a significance threshold alpha of 0.05 were designated as artefactual and rejected from further analysis. Homogeneous field correction (HFC) was applied via MNE-Python to reduce interference that manifests as a homogeneous field in the MSR (Seymour et al., 2022; Tierney, Alexander, et al., 2021). Continuous data were bandpass filtered (2-50 Hz using a 4th-order IIR Butterworth filter). The data then underwent Independent Component Analysis (ICA). The fastICA algorithm (Hyvarinen, 1999) was applied to these segmented data as implemented in MNE Python. Components containing cardiac artefacts and eyeblinks (identified in time course and topographies of the ICA components) were removed for each subject.

### 2.6 Coregistration & Forward Modelling

To facilitate accurate source reconstruction and localisation, a coregistration and 3D digitisation procedure was performed to register the OPM sensor positions and orientations to the participant’s specific head anatomy (Rhodes et al., 2025). 3D digitisation was conducted using an EinScan handheld scanner (EinScan H, Version 1.2.1.1, SHINING 3D, Hangzhou, China, support.einscan.com), which uses an infrared Class I laser light and white LEDs to generate a high-resolution 3D surface image. The scanner projects a known pattern of light detected by a camera. The camera captured many such frames of data per second, showing how different patterns of light are distorted, and by analysing these data it constructs point clouds depicting the 3D surface of the head. This technique is suitable for use with participants who struggle to remain still and is thought to be more practical and cost-effective than MRI (Rhodes et al., 2025). A scan of each individual wearing the helmet was combined with geometry of the helmet (using a CAD image) to delineate the sensor locations and orientations relative to the face and scalp using the Cerca Magnetics cREG and cFIFF software. This then produced a file containing MEG signals, as well as events and coregistration information all simultaneously stored in the BIDS data format. Additionally, six subjects underwent a T1 weighted anatomical MRI scan (3T Siemens; MPRAGE sequence; 1 mm isotropic resolution). By extracting the scalp surface from the MRI using osl-ephys and FSL-BET (Quinn et al., 2022), we then fit the extracted surface to the optical digitisation of the head (without the helmet). This enabled complete coregistration of the sensor locations/orientations to brain anatomy (Rivero et al., 2025). Any headshape points skewing the coregistration could be deleted using the custom RHINO function delete_headshape_points (van Es et al., 2025).

For the use of template-based MRIs, we utilised the same procedure, but instead using RHINO (Quinn et al., 2022) to warp the MNI152 template (Evans et al., 2012) to the participant’s head shape digitised via the EinScan. To do so, coregistration between the head model and the OPM-MEG geometry was refined using an iterative closest point (ICP) procedure using a full affine (12-parameter) transformation. The scalp surface extracted from the template (FSL BET outer-skin surface) and the digitised head-surface points were first expressed in a common coordinate frame. The digitised points were downsampled onto a 1-cm grid by spatial averaging; points more than 2 cm below the nasion, together with facial points, were discarded so that fitting was driven by the cranial surface. At each iteration, every headshape point was matched to its nearest scalp-mesh vertex, correspondences separated by more than 32 mm were rejected, and a 12-parameter affine transform (three translations, three rotations, three anisotropic scalings and three shears) was estimated in closed form by linear least squares (Moore-Penrose pseudoinverse). Iterations continued until convergence or a maximum of 128 iterations. The resulting affine was then applied to deform the outer-skin and inner-skull surface meshes so that the head model conformed to the measured headshape, and these warped surfaces were used for subsequent forward modelling. Following successful coregistration, a single-shell boundary element model (BEM) was constructed based on the brain surface extracted (Dale et al., 1999; Fischl, 2012). This model was subsequently used to generate a volumetric forward model on a 5-mm grid encompassing the entire brain volume (the two models can be viewed in the supplementary materials for both pipelines). The lead field matrix was then computed according to the participant’s head position relative to the OPM sensor array. The code for this is provided alongside the data.

### 2.7 Data Analysis of the Theta Band (4-8 Hz)

The 4-8 Hz band was defined a priori to match previous conventional and OPM-MEG studies using a similar paradigm (Barry, Barnes, et al., 2019; Barry, Tierney, et al., 2019; Tierney, Levy, et al., 2021), as well as the typical theta range used in the majority of neuroimaging studies focused on the hippocampus and memory (Herweg et al., 2020). Dynamic Imaging of Coherent Sources (DICS) was used to localise modulations in oscillatory brain activity (Gross et al., 2001). To identify differences in the theta band, cross-spectral density (CSD) was computed for the 4-8 Hz range using a multi-taper method with 2 Hz spectral smoothing. For the analysis of theta power, CSD matrices were calculated for the same trial interval: 0ms - 3000ms. These CSDs were combined and used with the forward model to construct a common spatial filter. The data rank was estimated first, and for each source, the orientation was optimised to maximise the output power. For depth-weighting, the default setting for MNE python was used (depth = 1). It has been shown previously that deep-sources elicit the largest amount of error during source reconstruction in MEG (Bénar et al., 2021). For example, a depth-weighting adjustment of 1 during beamforming is most useful for deeper sources, with superficial sources benefitting most from a depth-weighting of 0 (Afnan et al., 2026; Bénar et al., 2021; Pascarella et al., 2023). In addition, we only compute a condition-based contrast using the same spatial filter. The computed common spatial filter was then applied to the condition-specific CSDs to compute the decibel difference in theta power between scene and counting trials, using a regularisation parameter of 5% (Brookes et al., 2008). Therefore, any depth-weighting bias would scale across conditions equally. Remaining depth-related limitations, namely lower signal-to-noise ratio and broader spatial spread for deep sources, are considered later in the Discussion. For analysis, the resulting power change map was transformed to Montreal Neurological Institute (MNI) space using the transformation estimated from the registration of the native T1-weighted MRI to the MNI template sampled on a 5-mm grid. For the template pipeline, all power maps were already in MNI space. The resulting images were smoothed using a 9-mm Gaussian kernel.

## 3. Results

### 3.1 Behaviour

Participants completed on average (Mean ± SD) 93.5 (± 23.1) scene and 78.9 (± 19.9 SD) counting trials. Retention after exclusion of timeouts, missed responses, and any unsuccessful trials resulted in an average of 94.1% successful scene trials (± 4.0%); and 96.3% successful counting trials (± 3.2%). For the number trials analysed per condition for each participant, see supplementary materials. Since performance could not be measured objectively on both imagination and mentally counting, participants were required to provide a subjective judgment of imagination and counting success. Therefore, reaction times (length of time it took participants to provide feedback following trial completion) were measured as a proxy for task attention. Using a paired *t*-test we found that reaction times did not differ between the scene (0.995 seconds ± 0.307 seconds) and counting (1.02 seconds ± 0.274 seconds) conditions (*t*(14) = - 0.894, *p* = 0.387, Cohen’s *d* = 0.231).

### 3.2 Theta power attenuation during scene imagination without an MRI

To confirm that the 12-parameter template warping pipeline recovers consistent medial temporal lobe and possibly hippocampal signatures in OPM-MEG at the group level in a larger sample (n = 15), we calculated the difference in theta (4-8 Hz) power between scene imagination and counting baseline trials. Following coregistration, we then ran surface extraction and forward modelling within the template pipeline. At a whole brain level, theta power was attenuated during novel scene imagination compared to counting (mean = -0.198 dB ± 0.434 dB; Figure 2a-c). The global mean attenuation peaked at -0.699 dB located at MNI (x = -35, y = -6, z = -12), labelled as the left insula by the AAL3 atlas (Rolls et al., 2020). To statistically investigate this activity, we used a whole-brain cluster-based permutation *t*-test (one-sample, two-tailed) on the scene-versus-counting power difference across participants (5000 permutations; cluster-forming threshold |*t*| = 3.33, corresponding to a two-tailed *p* = 0.005). This identified a single significant cluster (109 voxels, *p* = 0.048) of negative theta power change during scene imagination relative to counting, peaking in the left parahippocampal gyrus (MNI x = -15, y = -26, z = -17, *t* = -3.55) with its centre of mass in the left hippocampus (*x* = -17, y = -26, z = -16). Of the 55 voxels labelled in the AAL3 atlas, 82% (45/55) fell within the left medial temporal lobe; 47% (26) in parahippocampal gyrus and 35% (19) in hippocampus. No voxels fell within the lateral or ventral temporal neocortex. The remaining ten labelled voxels were isolated (≤2 each) across left cerebellar, fusiform, thalamic and midbrain regions. No other cluster approached significance (next largest *p* = 0.17). At a more lenient cluster-forming threshold (|*t*| = 2.98, two-tailed *p* = 0.01) the same effect formed a larger cluster (583 voxels, *p* = 0.033) with an identical peak, extending further into lateral and ventral temporal cortex; peak location, sign and hemisphere were unchanged across thresholds.

**Figure 2:**
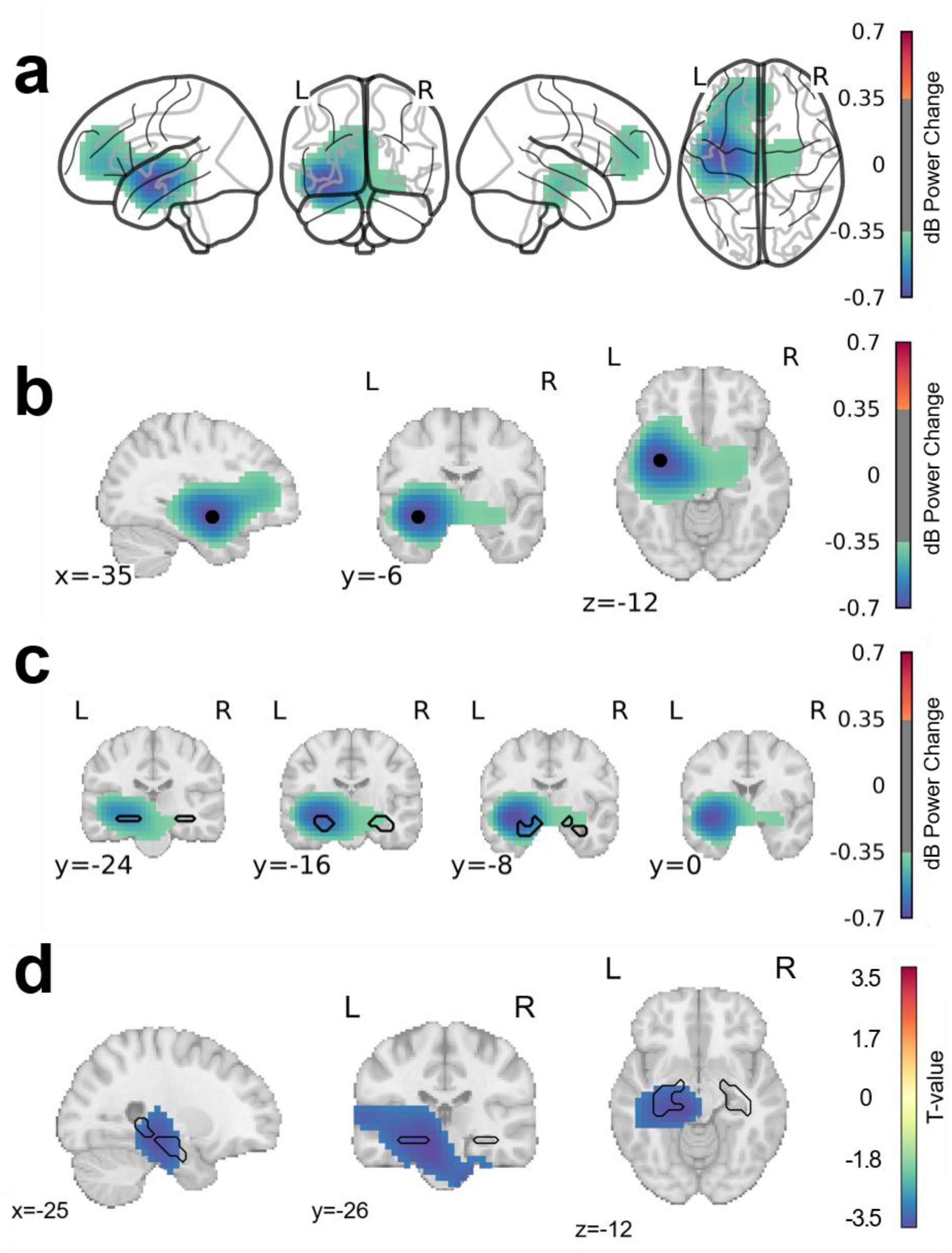
Group-level source reconstruction of scene vs counting using template pipeline. Source reconstruction of the scene-versus-counting theta (4-8 Hz) contrast across the full sample, computed with the template (MNI152) coregistration and forward-model pipeline. (a) **Whole-brain theta power change map**; attenuation during scene imagination peaked at MNI x = -35, y = -6, z = -12 (−0.699 dB). **(b) Orthogonal slices centred on the group global attenuation peak** (sagittal x = -35; coronal y = -6; axial z = -12), reconstructed with the template pipeline. Colour denotes theta (4-8 Hz) power change during scene imagination relative to counting, in decibels. Attenuation was maximal in the left anterior medial temporal lobe, encompassing the left hippocampus and adjacent parahippocampal and temporal cortex. (c) **Coronal slices (y = -24, -16, -8, 0) through the medial temporal lobe**, with the left and right hippocampus outlined in black (AAL3 atlas). Theta attenuation overlapped the left hippocampal region and extended into neighbouring parahippocampal and temporal cortex. Maps are displayed with a symmetric colour scale (±0.70 dB) and thresholded at ±0.35 dB for visualisation, such that voxels between -0.35 and +0.35 dB are not shown. This is for visualisation purposes and does not denote statistical significance. (d) **Results of cluster-based permutation t-test.** The single significant cluster from a cluster-based permutation *t*-test (one-sample, two-tailed; 5000 permutations; cluster-forming threshold *t* = 3.33, *p =* 0.005), peaking in the left parahippocampal gyrus (x = -15, y = -26, z = -17, *t* = -3.55, *p* = 0.048). The above figure is orthogonally sliced at the centroid of the significant cluster (x = -17, y = -26, z = -16). Colour bar denotes *t*-values and are not thresholded; only significant voxels are shown, with negative values indicating significant (*p* < 0.05) theta attenuation. L = left hemisphere and R = right hemisphere.

### 3.3 MRI-free OPM-MEG source reconstruction alignment

The setup for our current study provided us with a useful opportunity to compare the accuracy of using T1 MRI scans against a readily available template in the FSL package (MNI152) for source reconstruction (including the calculation of an individual’s forward model). In order to explore this, we isolated participants who had undergone a T1 MRI (n=6). Comparing the two pipelines, we showed theta attenuation across the whole brain in both models (MRI whole-brain mean = -0.036 ± 0.081 dB; Template = -0.045 ± 0.097 dB; individual whole-brain activity from all participants from both pipelines is available in the supplementary material). Masking the hippocampus bilaterally using the AAL3 atlas (Rolls et al., 2020), theta attenuation was substantially stronger within the hippocampal ROI in both reconstructions (MRI = -0.297 ± 0.193 dB; Template = -0.325 ± 0.218 dB). The peak hippocampal effect localised to (x = -25, y = -11, z = -17; -0.599 dB) in the MRI-based solution and to (x = -30, y = -6, z = -17; -0.698 dB) in the template-based solution. This is a separation of ∼7 mm. The unconstrained whole-brain peak coincided with the hippocampal peak in the MRI-based solution (Figure 3a), whereas in the template-based solution it fell in the adjacent left superior temporal pole (x = -40, y = -1, z = -22; -0.721 dB; Figure 3b). Given the 5 mm grid resolution and the spatial spread of the beamformer estimate, a displacement of this order is within the localisation uncertainty expected from coregistration error and helmet movement.

**Figure 3:**
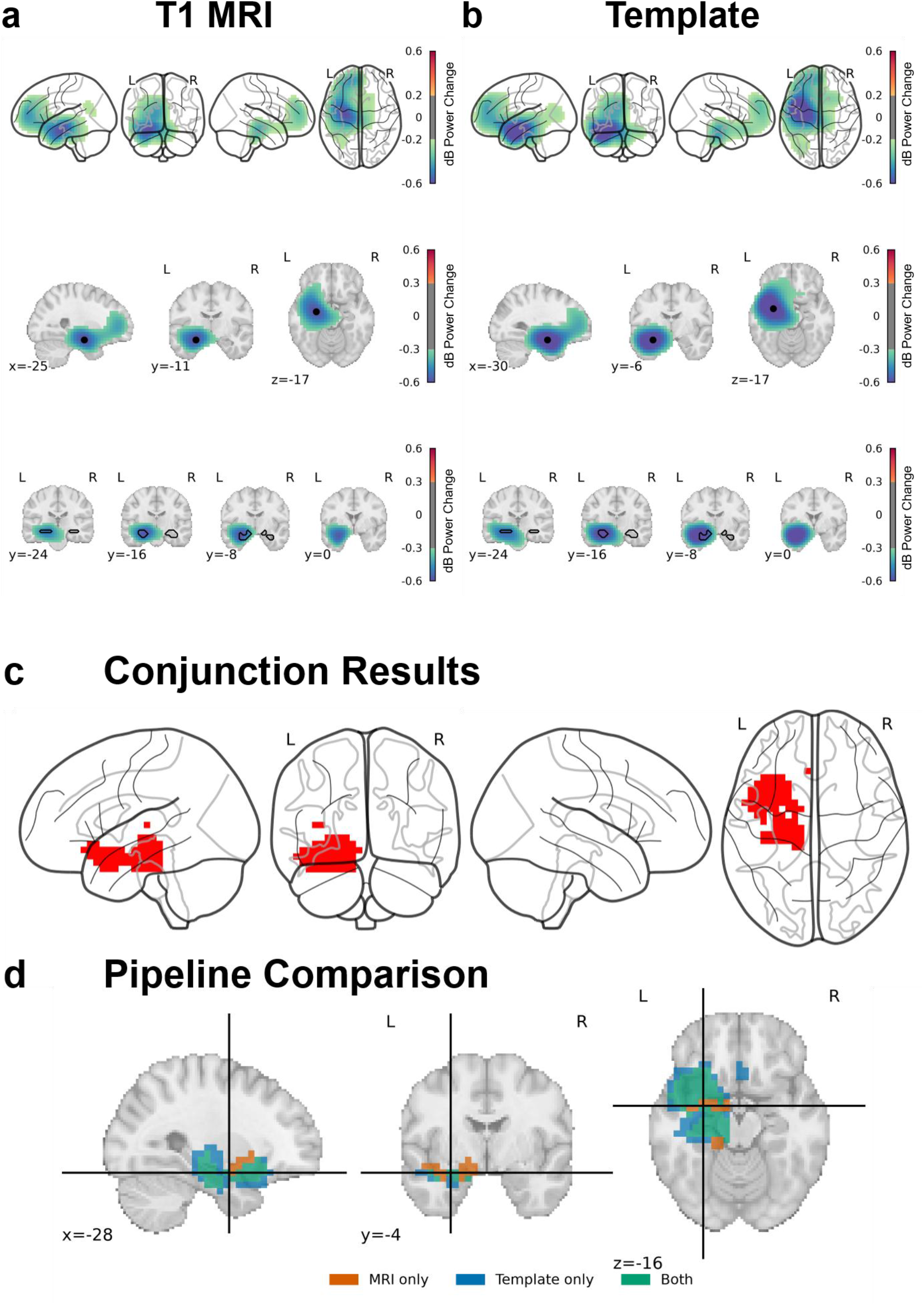
Source reconstruction pipeline comparisons of global peaks and results of conjunction analyses for T1 MRI and Template MRI. Group-average theta power change (dB) for the scene-versus-number contrast, reconstructed using (a) individual T1-derived head models and (b) template models. In each panel the top row shows a whole-brain glass-brain projection (thresholded at a lower value to display the full spatial extent of the effect), the middle row an orthogonal view centred on the peak hippocampal voxel (black circle), and the bottom row a posterior-to-anterior coronal montage (y = -24 to 0 mm) through the medial temporal lobe, with the AAL3-defined hippocampus outlined in black. The peak effect localised to the left hippocampus in both models, at (x = -25, y = -11, z = -17) for the T1 and (x = -30, y = -6, z = -17) for the template reconstruction, a separation of approximately one voxel at the 5 mm grid resolution. (c) Conjunction maps showing voxels at which the effect was consistent across 5/6 participants, for the T1 and template displayed as glass-brain projections. Orthogonal slices through the conjunction centroid (d), showing consistent voxels for between-subjects MRI (orange), Template (blue) and between-pipeline voxels (green). Colour bars denote theta power change in decibels; and are thresholded for visualisation purposes only with 33% of the maximum power for glass brain plots, and 50% of max power for hippocampal and temporal lobe comparisons.

To assess whether the two reconstruction pipelines localised consistently, we quantified their spatial overlap using the Dice similarity coefficient (DSC), which has been used previously in MRI and MEG research (Dice, 1945; Miyazaki et al., 2026; Zou et al., 2004). Each subject’s contrast was binarised at the 95th percentile of its own distribution within each subject’s beamformer contrast before overlap was computed. Across the six subjects, the per-subject MRI-versus-template overlap was consistent: Dice = 0.74 ± 0.05 (mean ± SD), with per-subject Dice ranging from 0.69 to 0.83. We then ran a group-level conjunction analysis for each pipeline, retaining voxels active in at least five of the six subjects. At the strictest conjunction (only showing voxels in all six subjects) the T1 MRI pipeline retained 23 voxels, whereas the template pipeline retained only 2; centroid comparison at this level was therefore not meaningful. We report the ≥5/6 conjunction, where both pipelines yield robust voxel counts, as the basis for spatial comparison. This yielded 302 consistent voxels for the MRI pipeline and 419 for the template pipeline, with a Dice coefficient of 0.59. The centroids of these conjunction maps lay at MNI (x = -27.6, y = -4.1, z = -14.9) for the T1-MRI and (x = -30.4, y = -5.2, z = -15.0) for the template. These yield a Euclidean separation of 3.0 mm. This is sub-voxel spatial accuracy as it is below our 5 mm grid resolution (0.6 voxels). Both pipelines therefore localised to the left temporal lobe with near-identical centres of mass, indicating that template-based pipeline reproduced similar results to the T1-MRI pipeline to within a single voxel in at least 5 subjects. Correlation analyses similar to Rhodes et al. (2025) of task-relevant theta power recorded from anatomical regions using the MRI and Template pipelines are provided in the supplementary materials.

## 4. Discussion

In the current study, we successfully deployed OPM-MEG to measure brain activity during a hippocampal-dependent task. We demonstrated that source reconstruction using a template MRI for coregistration yielded patterns of activity closely comparable to those obtained with an individual structural MRI, and that this template pipeline recovered the previously reported theta effect associated with the scene-construction network at the group level. Across fifteen adults, novel scene imagination was accompanied by a significant attenuation of theta (4-8 Hz) power relative to a counting baseline, forming a single cluster that peaked in the left parahippocampal gyrus with its centre of mass in the left hippocampus and the majority of its labelled voxels confined to the left medial temporal lobe. Contrary to our a priori hypothesis, this effect was left-lateralised.

### 4.1 Deep medial temporal lobe effects are recoverable without an MRI

We successfully reported a significant cluster of activity centred around the medial temporal lobe during scene imagination, which, similar to previous research, represented a decrease below the baseline counting condition (Barry, Barnes, et al., 2019; Barry, Tierney, et al., 2019; Lucie-Read et al., 2024). Most importantly, this was recoverable without an individual MRI scan. We ensured that source localisation was optimised for the MTL by increasing sensor number, using triaxial measurement, strategically placing sensors near the back and sides of the head, as well as increasing the number of trials (Quraan et al., 2011). We additionally extend the work of Rhodes et al. (2025) by demonstrating that similar template-modelling of source activity can extend beyond cortical surfaces, and recover deeper, more challenging sources if the setup and paradigm are appropriate. Nevertheless, we report larger separations between the pipelines. This is, to our knowledge, the first demonstration that a deep-source task-related oscillatory effect can be reconstructed at the group level using OPM-MEG without the requirement for individual anatomy. We leveraged a hippocampal-dependent task for the current study, and it is likely that should investigation of superficial sources be exploratory in nature, recovering deeper activity may prove problematic (Jaiswal et al., 2025). We would suggest that increasing trial numbers and participants should alleviate these issues. One suggestion by Rhodes et al. was that it is likely that differences between T1 MRI and warping are random and will average out at the group level. We would further support this interpretation and additionally add that warping of a structured light scan to a template readily available in the FSL package (such as MNI152 used here) may be a viable alternative. Therefore, we have made our 12-parameter warping code and pipeline available to the OPM-MEG community (https://doi.org/10.5281/zenodo.21721282). Considering the larger errors reported in the localisations of peak activity, we would strongly recommend that template reconstruction for precise source analysis (for example, in epilepsy seizure localisation) be avoided. It is likely that the MRI results are better localised and may be better at localising the centre of mass of significant activity, if clinically relevant. However, whilst a structural MRI is the gold standard, for cognitive neuroscience research with OPMs, it is likely not necessary. The whole-brain centroid of activity from our conjunctions was separated by approximately 3 mm and we could successfully recover our expected effects with statistical significance. Importantly, the use of light-scanned head meshes allowing MRI templates to be warped to head shape is essential. We would envisage that without this resolution, localising to these areas would prove difficult (Hill et al., 2020). However, this may be something that different research labs with different setups could consider examining.

### 4.2 Scene imagination as a functional benchmark for subcortical OPM-MEG

Although a power decrease may appear counterintuitive for a region engaged in scene construction, MTL theta decreases are commonly reported in intracranial studies (Herweg et al., 2020). These decreases have been proposed to reflect broadband changes associated with task engagement, which can mask more localised memory-related theta increases. Because scene imagination and counting differ substantially in their cognitive demands, the decreases observed here may partly reflect these differences. Future work could examine whether theta-band connectivity between the MTL and other regions differs between conditions. Nevertheless, the theta attenuation we observed replicates a previously reported signature of the scene-construction network and is consistent with reports of medial temporal theta decreases (Barry, Barnes, et al., 2019; Lucie-Read et al., 2024). Within this account, novel scene imagination may capture the formation of an early spatial schematic, generated by the hippocampus together with the ventromedial prefrontal cortex, that then scaffolds subsequent episodic memory. Our data demonstrate that the previously demonstrated signature of this process is recoverable at the group level with wearable OPM-MEG without the need for individual anatomy. This may be a useful task to use as a benchmark in OPM-MEG, to initially accurately reconstruct each subjects MTL & hippocampal activity prior to testing other, more naturalistic tasks. In rodents, hippocampal theta is classically a movement and navigation-related rhythm whose power increases during active exploration (Buzsáki & Moser, 2013; Herweg et al., 2020). It is possible, that the attenuation reported here and in previous work (Barry, Barnes, et al., 2019; Barry, Tierney, et al., 2019; Dalton et al., 2018; Lucie-Read et al., 2024; Zeidman et al., 2015) could be a signature of the hippocampus binding cortical inputs into a coherent spatial framework - the operation that scene construction is proposed to instantiate (Zeidman & Maguire, 2016). However, we do not directly test for that here. It is worth considering that this effect is inherently relative to our baseline. Because we contrast conditions, and verbal working memory is itself associated with frontal-midline theta power (Ishii et al., 2014), it is possible that our observed attenuation may reflect greater theta engagement during counting rather than suppression during imagination. Future studies may employ different baseline comparisons. However, as argued by Barry et al., using a resting baseline has been shown to involve recurrent and spontaneous hippocampal activity (from the default mode network) in both fMRI and MEG (Barry, Barnes, et al., 2019; Fox & Raichle, 2007; Raichle, 2015; Vincent et al., 2006). Additionally, the pre-beep baseline contains eyes-open data, compared to the task-related data, which is eyes closed. This inherently makes analysis within-condition inherently difficult. Whilst connectivity analyses may help investigate this, our data were not sufficiently powered to provide any insight, and it is beyond the scope of this paper. Nevertheless, OPMs’ tolerance for longer, more naturalistic recordings are well placed to investigate this further.

### 4.3 Spatial resolution and anatomical precision at depth

Some caution is warranted in attributing this effect to the hippocampus in isolation. The significant cluster peaked in the parahippocampal gyrus, although its centre of mass fell within the hippocampus, parahippocampal voxels outnumbered hippocampal (26 vs 19). The beamformer resolution at depth cannot cleanly separate the two. Our results therefore localise the effect to the left medial temporal lobe rather than to the hippocampus *specifically*. Whilst our a priori hypothesis focused on the medial temporal lobe, specifically the hippocampus, we do observe activity from several other places from our group-level average, that did not reach significance. One well supported area involved in scene reconstruction is the ventromedial prefrontal cortex (vmPFC). We do see some frontal activity at the group level at the cortex. Whilst this did not survive our cluster permutation test, there is also the possibility that the counting task used as a baseline, may have activated this region, which was then undetectable during the calculation of the difference between these conditions. We have provided the group unthresholded map on Zenodo for hypothesis generation.

Agreement between MRI and template pipelines show that the template introduced little additional error (Rhodes et al., 2025), but not that either pipeline localised the effect accurately (i.e., there is no ground-truth). Localisation error increases with source depth, and as previously mentioned, validation studies with simultaneous intracranial recordings show that optimal source-imaging parameters depend on depth (Afnan et al., 2026; Bénar et al., 2021). Our attribution of the effect to the left MTL should therefore be read as approximate.

Furthermore, contrary to our hypothesis, we reported a significant theta attenuation in the left medial temporal lobe as well as the left hippocampus, with the peak in the left parahippocampal gyrus. Our original hypothesis was that we would observe this effect in the right hippocampus, as this was reported in OPMs by Barry et al. (2019). However, it is perhaps not surprising that this hypothesis was unsupported. We would suggest that it is likely that by increasing trial count, sensor positioning, axial measurements and the number of sensors alongside increasing the sample size; we are closer to the increased resolution provided by conventional MEG. In these studies, we see a clear effect localised to the left hippocampal region (Barry, Barnes, et al., 2019; Lucie-Read et al., 2024). This provides us with further confidence in our current results that we have replicated an effect typically found using conventional MEG, using OPMs combined with a template MRI. Nevertheless, we could not rule out that right hippocampal engagement is significantly contributing to this network and is simply undetectable by our beamforming analysis (Pu et al., 2018; Ruzich et al., 2019).

A concern for any study of deep sources using MEG, is that activity attributed to deeper sources may partly reflect leakage from cortical regions and other frequency bands, such as occipital alpha leaking into our theta activity in the MTL (see Ruzich et al., 2019). We consider this an unlikely explanation for our current findings. Firstly, both conditions were performed with eyes closed, so any eyes-closed posterior alpha was matched across conditions and would cancel out in the contrast calculations. Secondly, Quraan et al. (2011) suggest that in order to recover hippocampal signals in conventional MEG, increased trials and group averaging is key with a control-condition subtraction acting as a natural signal leakage-reduction. Our study was designed to minimise leakage, by computing a common spatial filter, applying it to each condition and reporting power from the condition contrast. In addition, the significant cluster contained no occipital or parietal voxels at all. Over 82% of its labelled voxels lay in the left MTL, with the remainder scattered across cerebellar, fusiform, thalamic and midbrain regions (Section 3.2). Finally, the direction and location of our effect is consistent with previous conventional MEG work from Barry et al. (2019) using the same task. Repeating the analysis in the alpha band (8-12 Hz) revealed no significant clusters (supplementary materials). With the closer distance of the OPMs to the scalp (approx. 1cm), we would argue that it is very likely we have recovered a true theta effect, centred around the left medial temporal lobe and hippocampus without the need for an MRI.

### 4.4 Limitations and future directions

There are several limitations to this study that must be acknowledged explicitly. Our main effect was derived from fifteen participants and the MRI-versus-template validation from six; while sufficient to demonstrate feasibility, these samples are modest for a methodological validation. The group effect discussed here comes from a single cluster that survived a stringent cluster-forming threshold (*p* = 0.048). This was larger and more significant at a more lenient threshold (*p* = 0.033). Whilst robust to permutation testing, we interpret our findings cautiously and do not over-extend claims related to the theoretical support of the scene-construction network itself. Our sample comprised young, healthy adults, and extension to the clinical populations that motivate this work remains to be demonstrated. Similarly to Rhodes et al., our template MRI was not sex-matched nor in this instance, age-matched.

Here, we demonstrated our method with an openly accessible and easy to use template available in a standard neuroimaging package such as FSL (Jenkinson et al., 2012; Laird et al., 2011). Being selective with templates for warping might improve results. Additionally, testing this template warping with patients, older adults or children would be required before attempting to explore any clinical or developmentally relevant questions (Jaiswal et al., 2025). Deploying this scene-imagination task with clinical groups is very feasible. However, we would suggest shorter blocks and fewer trials per participant to reduce fatigue. We would further suggest that researchers examine mental imagery ability using validated scales (Andrade et al., 2014). This ability is heavily related to autobiographical memory ability in Alzheimer’s Disease patients, a neurodegenerative disorder that disproportionately affects the hippocampus (El Haj et al., 2016, 2019). Because scene-construction success could not be measured objectively, we relied on subjective ratings as a marker of task engagement. Including a memory test afterwards outside the MSR (Barry, Barnes, et al., 2019), as well as screening for aphantasia would have been useful for understanding performance. Aphantasic individuals tend to have less, but similar patterns of hippocampal activation compared to controls (Monzel et al., 2024; Zeman et al., 2015). Whilst these studies use fMRI and do not make claims about oscillations, it nonetheless could provide greater confidence in our results.

A further consideration is that our effects were computed as a scene vs counting contrast, so a genuine theta decrease during scene imagination and an increase during counting cannot be distinguished; the attenuation we report should be interpreted relatively. Mental counting has been shown to increase frontal-midline theta oscillations (Gärtner et al., 2015; Ishii et al., 2014). Whilst our significant cluster was confined to the left MTL but cannot exclusively rule out any counting-related theta increases within the MTL itself. Incorporating a closed eyes resting period prior to imagination or counting may help resolve this. Our left-lateralised result is consistent with the broader fMRI and SQUID-MEG literature and the discrepancy to the findings reported by Barry et al. may reflect differences in sample size and sensor coverage rather than a genuine reversal of lateralisation. Future work should attempt extending this approach to naturalistic and virtual-reality paradigms (Zabbah et al., 2026). Furthermore, whilst several features of our data argue against alpha leakage, we cannot exclude it entirely. Contrasting against a control condition reduces but does not eliminate leakage into deep structures (Quraan et al., 2011). It is possible that activity at the 8 Hz boundary, along with individual alpha frequency variation suggests that the theta band could include some low-frequency alpha activity from cortical sources. Future work should quantify this by controlling for eye condition or using concurrent intracranial recordings combined with a hippocampal dependent task such as ours. All data are available via Zenodo (https://zenodo.org/records/21721282). We actively encourage hypothesis generation and/or further research using these data. This is useful should researchers also wish to take a data-driven approach to understanding neural activity during scene imagination, which was outside the scope of this manuscript.

## 5. Conclusions

Using a triaxial 60-sensor OPM-MEG array and a scene imagination task, we recovered the previously reported theta attenuation of the scene-construction network at the group level, localised to the left medial temporal lobe with its peak in parahippocampal cortex and centre of mass in the left hippocampus. This effect was reconstructed using a headshape warping procedure and a template MRI. Together, these findings support the use of OPM-MEG to study medial temporal lobe function without individual structural imaging. This is important for deploying OPM-MEG with populations for whom MRI is difficult, reducing the cost and time of neuroimaging protocols and making cognitive neuroscience research accessible in labs that have OPM-MEG but lack the resources for structural scanning.

## Data and Code Availability

The data will be made available in BIDS format and the scripts as Jupyter Notebooks via Zenodo upon manuscript acceptance: https://doi.org/10.5281/zenodo.21721282. These data do not include the raw EinScan head scans but can be requested from the corresponding author upon reasonable request. The 12-parameter warping code and full pipeline are available alongside these data. For further analysis, we followed the OPM-FLUX pipeline, for which scripts are available here https://github.com/FLUX-pipeline/Cerca/. Some additional analysis scripts that rely on the use of OSL-ephys (RHINO/FSL) with tutorial & data accessible via https://github.com/neurofractal/OPM-oxford/tree/main/tutorials.

## Author Contributions

**Conor Thornberry**: Project Administration (lead); Funding Acquisition (lead); Conceptualization (lead); writing – original draft (lead); formal analysis (lead); writing – review and editing (equal); Methodology (lead); Investigation (lead); Software (equal). **Prathiksha Math**: Investigation (supporting); Writing – Review & Editing (supporting); Methodology (supporting). **Margarida Cohen Serra**: Investigation (supporting); Writing – Review & Editing (supporting); Methodology (supporting). **Robert Seymour**: Conceptualization (supporting); Writing – review and editing (equal); Software (equal). **Clodagh Nolan**: Investigation (supporting); Writing – Review & Editing (supporting); Methodology (supporting). **Robert Whelan**: Supervision (lead); Writing – review and editing (equal); Resources (lead).

## Acknowledgements

We would like to thank Sajjad Zabbah & Nic Alexander for providing the word list used in this task. We would also like to thank Lena Gacek and Katie Zorba for their help with early testing and setup of the OPM. We would like to thank Prof Redmond O’Connell for his encouragement and support with the use of the OPM at TCD. Thank you to the Royal Irish Academy for funding a visit to Cardiff University. As a result, thank you to Prof Krish Singh, Dr Haydee Garcia-Lazaro & Dr Luke Tait at Cardiff University for their time and detailed explanations of MEG analysis. This manuscript is inspired by and written in memory of Irish neuroscientist Prof Eleanor Maguire.

## References

1. Afnan, J., Fratello, M., Bonini, F., Medina Villalon, S., Cai, Z., Lina, J.-M., Badier, J.-M., Bartolomei, F., Gotman, J., Bénar, C. G., & Grova, C. (2026). Assessing Strengths and Limitations of Magnetoencephalography Source Imaging With Intracerebral EEG. Advanced Science, 13(58), e76365. 10.1002/advs.76365

2. Aggleton, J. P., & O’Mara, S. M. (2022). The anterior thalamic nuclei: Core components of a tripartite episodic memory system. Nature Reviews. Neuroscience, 23(8), 505–516. 10.1038/s41583-022-00591-8

3. Andrade, J., May, J., Deeprose, C., Baugh, S.-J., & Ganis, G. (2014). Assessing vividness of mental imagery: The Plymouth Sensory Imagery Questionnaire. British Journal of Psychology, 105(4), 547–563. 10.1111/bjop.12050

4. Apostolova, L. G., Dutton, R. A., Dinov, I. D., Hayashi, K. M., Toga, A. W., Cummings, J. L., & Thompson, P. M. (2006). Conversion of Mild Cognitive Impairment to Alzheimer Disease Predicted by Hippocampal Atrophy Maps. Archives of Neurology, 63(5), 693–699. 10.1001/archneur.63.5.693

5. Aumont, E., Bedard, M.-A., Bussy, A., Arias, J. F., Tissot, C., Hall, B. J., Therriault, J., Rahmouni, N., Stevenson, J., Servaes, S., Macedo, A. C., Vitali, P., Poltronetti, N. M., Fliaguine, O., Trudel, L., Gauthier, S., Chakravarty, M. M., & Rosa-Neto, P. (2025). Hippocampal atrophy over two years in relation to tau, amyloid-β and memory in older adults. Neurobiology of Aging, 146, 48–57. 10.1016/j.neurobiolaging.2024.11.007

6. Barry, D. N., Barnes, G. R., Clark, I. A., & Maguire, E. A. (2019). The Neural Dynamics of Novel Scene Imagery. The Journal of Neuroscience, 39(22), 4375–4386. 10.1523/JNEUROSCI.2497-18.2019

7. Barry, D. N., Tierney, T. M., Holmes, N., Boto, E., Roberts, G., Leggett, J., Bowtell, R., Brookes, M. J., Barnes, G. R., & Maguire, E. A. (2019). Imaging the human hippocampus with optically-pumped magnetoencephalography. NeuroImage, 203, 116192. 10.1016/j.neuroimage.2019.116192

8. Bénar, C.-G., Velmurugan, J., López-Madrona, V. J., Pizzo, F., & Badier, J.-M. (2021). Detection and localization of deep sources in magnetoencephalography: A review. Current Opinion in Biomedical Engineering, 18, 100285. 10.1016/j.cobme.2021.100285

9. Bertossi, E., Aleo, F., Braghittoni, D., & Ciaramelli, E. (2016). Stuck in the here and now: Construction of fictitious and future experiences following ventromedial prefrontal damage. Neuropsychologia, 81, 107–116. 10.1016/j.neuropsychologia.2015.12.015

10. Boto, E., Holmes, N., Leggett, J., Roberts, G., Shah, V., Meyer, S. S., Muñoz, L. D., Mullinger, K. J., Tierney, T. M., Bestmann, S., Barnes, G. R., Bowtell, R., & Brookes, M. J. (2018). Moving magnetoencephalography towards real-world applications with a wearable system. Nature, 555(7698), 657–661. 10.1038/nature26147

11. Boto, E., Shah, V., Hill, R. M., Rhodes, N., Osborne, J., Doyle, C., Holmes, N., Rea, M., Leggett, J., Bowtell, R., & Brookes, M. J. (2022). Triaxial detection of the neuromagnetic field using optically-pumped magnetometry: Feasibility and application in children. NeuroImage, 252, 119027. 10.1016/j.neuroimage.2022.119027

12. Brookes, M. J., Leggett, J., Rea, M., Hill, R. M., Holmes, N., Boto, E., & Bowtell, R. (2022). Magnetoencephalography with optically pumped magnetometers (OPM-MEG): The next generation of functional neuroimaging. Trends in Neurosciences, 45(8), 621–634. 10.1016/j.tins.2022.05.008

13. Brookes, M. J., Vrba, J., Robinson, S. E., Stevenson, C. M., Peters, A. M., Barnes, G. R., Hillebrand, A., & Morris, P. G. (2008). Optimising experimental design for MEG beamformer imaging. NeuroImage, 39(4), 1788–1802. 10.1016/j.neuroimage.2007.09.050

14. Buzsáki, G., & Moser, E. I. (2013). Memory, navigation and theta rhythm in the hippocampal-entorhinal system. Nature Neuroscience, 16(2), 130–138. 10.1038/nn.3304

15. Chersi, F., & Burgess, N. (2015). The Cognitive Architecture of Spatial Navigation: Hippocampal and Striatal Contributions. Neuron, 88(1), 64–77. 10.1016/j.neuron.2015.09.021

16. Clark, I. A., Kim, M., & Maguire, E. A. (2018). Verbal Paired Associates and the Hippocampus: The Role of Scenes. Journal of Cognitive Neuroscience, 30(12), 1821–1845. 10.1162/jocn_a_01315

17. Dale, A. M., Fischl, B., & Sereno, M. I. (1999). Cortical Surface-Based Analysis: I. Segmentation and Surface Reconstruction. NeuroImage, 9(2), 179–194. 10.1006/nimg.1998.0395

18. Dalton, M. A., Zeidman, P., McCormick, C., & Maguire, E. A. (2018). Differentiable Processing of Objects, Associations, and Scenes within the Hippocampus. Journal of Neuroscience, 38(38), 8146–8159. 10.1523/JNEUROSCI.0263-18.2018

19. Dice, L. R. (1945). Measures of the Amount of Ecologic Association Between Species. Ecology, 26(3), 297–302. 10.2307/1932409

20. Ekstrom, A. D., & Hill, P. F. (2023). Spatial navigation and memory: A review of the similarities and differences relevant to brain models and age. Neuron, 111(7), 1037–1049.

21. El Haj, M., Gallouj, K., & Antoine, P. (2019). Mental imagery and autobiographical memory in Alzheimer’s disease. Neuropsychology, 33(5), 609–616. 10.1037/neu0000521

22. El Haj, M., Kapogiannis, D., & Antoine, P. (2016). Phenomenological Reliving and Visual Imagery During Autobiographical Recall in Alzheimer’s Disease. Journal of Alzheimer’s Disease, 52(2), 421–431. 10.3233/JAD-151122

23. Evans, A. C., Janke, A. L., Collins, D. L., & Baillet, S. (2012). Brain templates and atlases. NeuroImage, 20 YEARS OF fMRI, 62(2), 911–922. 10.1016/j.neuroimage.2012.01.024

24. Fischl, B. (2012). FreeSurfer. NeuroImage, 62(2), 774–781. 10.1016/j.neuroimage.2012.01.021

25. Fox, M. D., & Raichle, M. E. (2007). Spontaneous fluctuations in brain activity observed with functional magnetic resonance imaging. Nature Reviews Neuroscience, 8(9), 700–711. 10.1038/nrn2201

26. Gärtner, M., Grimm, S., & Bajbouj, M. (2015). Frontal midline theta oscillations during mental arithmetic: Effects of stress. Frontiers in Behavioral Neuroscience, 9, 96. 10.3389/fnbeh.2015.00096

27. Gohel, B., & Khare, M. (2023). EEG/MEG source imaging in the absence of subject’s brain MRI scan: Perspective on co-registration and MRI selection approach. International Journal of Imaging Systems and Technology, 33(1), 287–298. 10.1002/ima.22786

28. Gramfort, A., Luessi, M., Larson, E., Engemann, D. A., Strohmeier, D., Brodbeck, C., Goj, R., Jas, M., Brooks, T., Parkkonen, L., & Hämäläinen, M. (2013). MEG and EEG data analysis with MNE-Python. Frontiers in Neuroscience, 7. 10.3389/fnins.2013.00267

29. Greenberg, J. A., Burke, J. F., Haque, R., Kahana, M. J., & Zaghloul, K. A. (2015). Decreases in Theta and Increases in High Frequency Activity Underlie Associative Memory Encoding. NeuroImage, 114, 257–263. 10.1016/j.neuroimage.2015.03.077

30. Gross, J., Kujala, J., Hämäläinen, M., Timmermann, L., Schnitzler, A., & Salmelin, R. (2001). Dynamic imaging of coherent sources: Studying neural interactions in the human brain. Proceedings of the National Academy of Sciences, 98(2), 694–699. 10.1073/pnas.98.2.694

31. Hassabis, D., Kumaran, D., & Maguire, E. A. (2007). Using imagination to understand the neural basis of episodic memory. Journal of Neuroscience, 27(52), 14365– 14374. 10.1523/JNEUROSCI.4549-07.2007

32. Henson, R. N., Mattout, J., Phillips, C., & Friston, K. J. (2009). Selecting forward models for MEG source-reconstruction using model-evidence. NeuroImage, 46(1), 168–176. 10.1016/j.neuroimage.2009.01.062

33. Herweg, N. A., Solomon, E. A., & Kahana, M. J. (2020). Theta Oscillations in Human Memory. Trends in Cognitive Sciences, 24(3), 208–227. 10.1016/j.tics.2019.12.006

34. Hill, R. M., Boto, E., Rea, M., Holmes, N., Leggett, J., Coles, L. A., Papastavrou, M., Everton, S. K., Hunt, B. A. E., Sims, D., Osborne, J., Shah, V., Bowtell, R., & Brookes, M. J. (2020). Multi-channel whole-head OPM-MEG: Helmet design and a comparison with a conventional system. NeuroImage, 219, 116995. 10.1016/j.neuroimage.2020.116995

35. Holmes, N., Leggett, J., Boto, E., Roberts, G., Hill, R. M., Tierney, T. M., Shah, V., Barnes, G. R., Brookes, M. J., & Bowtell, R. (2018). A bi-planar coil system for nulling background magnetic fields in scalp mounted magnetoencephalography. NeuroImage, 181, 760–774. 10.1016/j.neuroimage.2018.07.028

36. Hyvarinen, A. (1999). Fast and robust fixed-point algorithms for independent component analysis. IEEE Transactions on Neural Networks, 10(3), 626–634. 10.1109/72.761722

37. Ishii, R., Canuet, L., Ishihara, T., Aoki, Y., Ikeda, S., Hata, M., Katsimichas, T., Gunji, A., Takahashi, H., Nakahachi, T., Iwase, M., & Takeda, M. (2014). Frontal midline theta rhythm and gamma power changes during focused attention on mental calculation: An MEG beamformer analysis. Frontiers in Human Neuroscience, 8. 10.3389/fnhum.2014.00406

38. Jaiswal, A., Nenonen, J., & Parkkonen, L. (2025). Pseudo-MRI Engine for MRI-Free Electromagnetic Source Imaging. Human Brain Mapping, 46(2), e70148. 10.1002/hbm.70148

39. Jenkinson, M., Beckmann, C. F., Behrens, T. E. J., Woolrich, M. W., & Smith, S. M. (2012). FSL. NeuroImage, 20 YEARS OF fMRI, 62(2), 782–790. 10.1016/j.neuroimage.2011.09.015

40. Laird, A. R., Eickhoff, S. B., Fox, P. M., Uecker, A. M., Ray, K. L., Saenz, J. J., McKay, D. R., Bzdok, D., Laird, R. W., Robinson, J. L., Turner, J. A., Turkeltaub, P. E., Lancaster, J. L., & Fox, P. T. (2011). The BrainMap strategy for standardization, sharing, and meta-analysis of neuroimaging data. BMC Research Notes, 4(1), 349. 10.1186/1756-0500-4-349

41. Long, N. M., Burke, J. F., & Kahana, M. J. (2014). Subsequent memory effect in intracranial and scalp EEG. NeuroImage, 84, 488–494. 10.1016/j.neuroimage.2013.08.052

42. Lucie-Read, M., Hodgetts, C. J., Lawrence, A. D., Evans, C. J., Singh, K. D., Umla-Runge, K., & Graham, K. S. (2024). Multimodal MEG and microstructure-MRI investigations of the human hippocampal scene network. Neuroscience. 10.1101/2024.07.16.603546

43. Maguire, E. A., Intraub, H., & Mullally, S. L. (2016). Scenes, Spaces, and Memory Traces. The Neuroscientist, 22(5), 432–439. 10.1177/1073858415600389

44. Maguire, E. A., & Mullally, S. L. (2013). The Hippocampus: A Manifesto for Change. Journal of Experimental Psychology. General, 142(4), 1180–1189. 10.1037/a0033650

45. Miyazaki, K., Nirasawa, S., Ishibashi, N., Akamatsu, K., Yamashita, O., & Miyawaki, Y. (2026). Suppression of information spreading in MEG source estimation using a functionally-structured Bayesian approach. NeuroImage, 333, 121934. 10.1016/j.neuroimage.2026.121934

46. Monk, A. M., Dalton, M. A., Barnes, G. R., & Maguire, E. A. (2021). The Role of Hippocampal-Ventromedial Prefrontal Cortex Neural Dynamics in Building Mental Representations. Journal of Cognitive Neuroscience, 33(1), 89–103. 10.1162/jocn_a_01634

47. Monzel, M., Leelaarporn, P., Lutz, T., Schultz, J., Brunheim, S., Reuter, M., & McCormick, C. (2024). Hippocampal-occipital connectivity reflects autobiographical memory deficits in aphantasia. eLife, 13, RP94916. 10.7554/eLife.94916

48. Pascarella, A., Mikulan, E., Sciacchitano, F., Sarasso, S., Rubino, A., Sartori, I., Cardinale, F., Zauli, F., Avanzini, P., Nobili, L., Pigorini, A., & Sorrentino, A. (2023). An *in–vivo* validation of ESI methods with focal sources. NeuroImage, 277, 120219. 10.1016/j.neuroimage.2023.120219

49. Pu, Y., Cheyne, D. O., Cornwell, B. R., & Johnson, B. W. (2018). Non-invasive Investigation of Human Hippocampal Rhythms Using Magnetoencephalography: A Review. Frontiers in Neuroscience, 12. 10.3389/fnins.2018.00273

50. Quinn, A. J., van Es, M. W. J., Gohil, C., & Woolrich, M. W. (2022). OHBA Software Library in Python (OSL). 10.5281/zenodo.6875060

51. Quraan, M. A., Moses, S. N., Hung, Y., Mills, T., & Taylor, M. J. (2011). Detection and localization of hippocampal activity using beamformers with MEG: A detailed investigation using simulations and empirical data. Human Brain Mapping, 32(5), 812–827. 10.1002/hbm.21068

52. Raichle, M. E. (2015). The Brain’s Default Mode Network. Annual Review of Neuroscience, 38(Volume 38, 2015), 433–447. 10.1146/annurev-neuro-071013-014030

53. Rakshit, A., Ghafari, T., Kowalczyk, A., & Jensen, O. (2026). OPM-FLUX: A Pipeline for OPM MEG Data Analysis (p. 2026.04.24.720604). bioRxiv. 10.64898/2026.04.24.720604

54. Rhodes, N., Rier, L., Boto, E., Hill, R. M., & Brookes, M. J. (2025). Source reconstruction without an MRI using optically pumped magnetometer-based magnetoencephalography. Imaging Neuroscience, 3, IMAG.a.8. 10.1162/IMAG.a.8

55. Rivero, G. R., Tanner, Z., Rier, L., Hill, R. M., Shah, V., Rea, M., Doyle, C., Osborne, J., Bobela, D., Morris, P. G., Mullinger, K. J., Boto, E., Holmes, N., & Brookes, M. J. (2025). OPM-MEG reveals dynamics of beta bursts underlying attentional processes in sensory cortex. Scientific Reports, 15(1), 30471. 10.1038/s41598-025-08037-8

56. Rolls, E. T., Huang, C.-C., Lin, C.-P., Feng, J., & Joliot, M. (2020). Automated anatomical labelling atlas 3. NeuroImage, 206, 116189. 10.1016/j.neuroimage.2019.116189

57. Ruzich, E., Crespo-García, M., Dalal, S. S., & Schneiderman, J. F. (2019). Characterizing hippocampal dynamics with MEG: A systematic review and evidence-based guidelines. Human Brain Mapping, 40(4), 1353–1375. 10.1002/hbm.24445

58. Schofield, H., Boto, E., Shah, V., Hill, R. M., Osborne, J., Rea, M., Doyle, C., Holmes, N., Bowtell, R., Woolger, D., & Brookes, M. J. (2022). Quantum enabled functional neuroimaging: The why and how of magnetoencephalography using optically pumped magnetometers. Contemporary Physics, 63(3), 161–179. 10.1080/00107514.2023.2182950

59. Schofield, H., Hill, R. M., Feys, O., Holmes, N., Osborne, J., Doyle, C., Bobela, D., Corvilain, P., Wens, V., Rier, L., Bowtell, R., Ferez, M., Mullinger, K. J., Coleman, S., Rhodes, N., Rea, M., Tanner, Z., Boto, E., de Tiège, X., … Brookes, M. J. (2024). A novel, robust, and portable platform for magnetoencephalography using optically-pumped magnetometers. Imaging Neuroscience, 2, imag–2–00283. 10.1162/imag_a_00283

60. Schofield, H., Hill, R. M., Rier, L., Kennett, E., Rivero, G. R., Gibson, J., Tyler, A., Tanner, Z., Worcester, F., Hayward, T., Osborne, J., Doyle, C., Shah, V., Boto, E., Holmes, N., & Brookes, M. J. (2025). Towards a 384-channel magnetoencephalography system based on optically pumped magnetometers. Imaging Neuroscience, 3, IMAG.a.1042. 10.1162/IMAG.a.1042

61. Seymour, R. A., Alexander, N., Mellor, S., O’Neill, G. C., Tierney, T. M., Barnes, G. R., & Maguire, E. A. (2022). Interference suppression techniques for OPM-based MEG: Opportunities and challenges. NeuroImage, 247, 118834. 10.1016/j.neuroimage.2021.118834

62. Thornberry, C., Caffrey, M., & Commins, S. (2023). Theta oscillatory power decreases in humans are associated with spatial learning in a virtual water maze task. European Journal of Neuroscience, 58(11), 4341–4356. 10.1111/ejn.16185

63. Tierney, T. M., Alexander, N., Mellor, S., Holmes, N., Seymour, R., O’Neill, G. C., Maguire, E. A., & Barnes, G. R. (2021). Modelling optically pumped magnetometer interference in MEG as a spatially homogeneous magnetic field. NeuroImage, 244, 118484. 10.1016/j.neuroimage.2021.118484

64. Tierney, T. M., Levy, A., Barry, D. N., Meyer, S. S., Shigihara, Y., Everatt, M., Mellor, S., Lopez, J. D., Bestmann, S., Holmes, N., Roberts, G., Hill, R. M., Boto, E., Leggett, J., Shah, V., Brookes, M. J., Bowtell, R., Maguire, E. A., & Barnes, G. R. (2021). Mouth magnetoencephalography: A unique perspective on the human hippocampus. Neuroimage, 225, 117443. 10.1016/j.neuroimage.2020.117443

65. van Es, M. W. J. W. J., Gohil, C., Quinn, A. J. J., & Woolrich, M. W. W. (2025). osl-ephys: A Python toolbox for the analysis of electrophysiology data. Frontiers in Neuroscience, 19. 10.3389/fnins.2025.1522675

66. Vincent, J. L., Snyder, A. Z., Fox, M. D., Shannon, B. J., Andrews, J. R., Raichle, M. E., & Buckner, R. L. (2006). Coherent Spontaneous Activity Identifies a Hippocampal-Parietal Memory Network. Journal of Neurophysiology, 96(6), 3517–3531. 10.1152/jn.00048.2006

67. Zabbah, S., Alexander, N. A., Mohammadi, Y., Mariola, A., Seymour, R. A., Puvvada, S., Barnes, G. R., & Bach, D. R. (2026). An integrated virtual reality platform for naturalistic neuroimaging with magnetoencephalography (p. 2026.01.28.701672). bioRxiv. 10.64898/2026.01.28.701672

68. Zeidman, P., & Maguire, E. A. (2016). Anterior hippocampus: The anatomy of perception, imagination and episodic memory. Nature Reviews Neuroscience, 17(3), 173–182. 10.1038/nrn.2015.24

69. Zeidman, P., Mullally, S. L., & Maguire, E. A. (2015). Constructing, Perceiving, and Maintaining Scenes: Hippocampal Activity and Connectivity. Cerebral Cortex, 25(10), 3836–3855. 10.1093/cercor/bhu266

70. Zeman, A., Dewar, M., & Della Sala, S. (2015). Lives without imagery – Congenital aphantasia. Cortex, 73, 378–380. 10.1016/j.cortex.2015.05.019

71. Zetter, R., Iivanainen, J., Stenroos, M., & Parkkonen, L. (2018). Requirements for Coregistration Accuracy in On-Scalp MEG. Brain Topography, 31(6), 931–948. 10.1007/s10548-018-0656-5

72. Zou, K. H., Warfield, S. K., Bharatha, A., Tempany, C. M. C., Kaus, M. R., Haker, S. J., Wells, W. M., Jolesz, F. A., & Kikinis, R. (2004). Statistical Validation of Image Segmentation Quality Based on a Spatial Overlap Index. Academic Radiology, 11(2), 178–189. 10.1016/S1076-6332(03)00671-8

