## supplementary materials for "MRI-free OPM-MEG recovers medial temporal lobe theta during scene imagination"

**Supplementary Figure 1: Forward model examples** from one participant (P001) for the MRI and Template coregistration and forward modelling solutions.

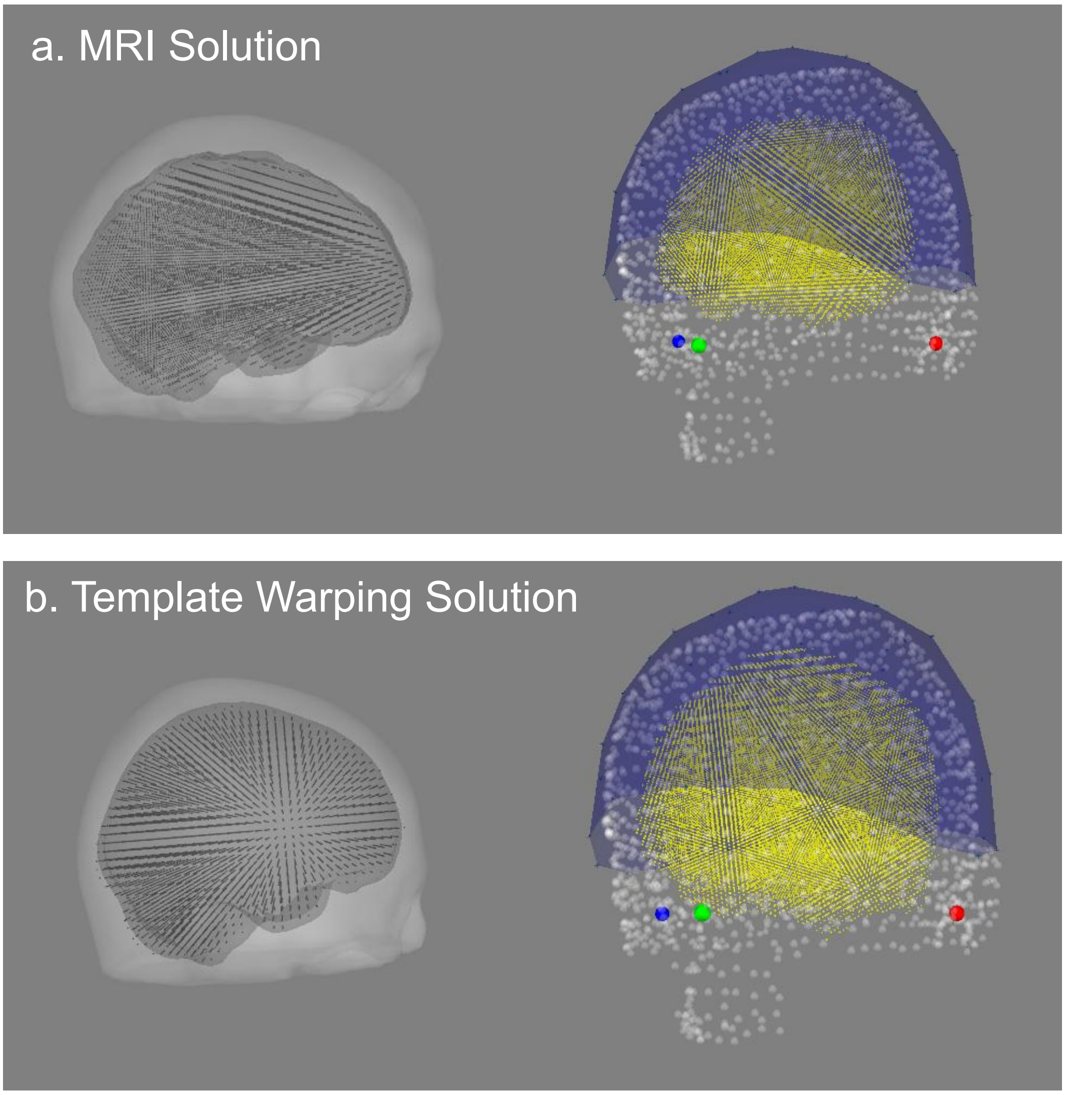

**Supplementary Table 1: Total trial count per participant.** Participant numbers correspond with BIDS data in Zenodo.

| *Participant* | *Trials analysed per condition* |
|---|---|
| 001 | 85 |
| 006 | 90 |
| 007 | 37 |
| 008 | 56 |
| 009 | 36 |
| 010 | 26 |
| 011 | 39 |
| 012 | 81 |
| 014 | 91 |
| 015 | 86 |
| 016 | 77 |
| 017 | 80 |
| 018 | 78 |
| 019 | 79 |
| 020 | 85 |

**Supplementary Figure 2: Individual subject glass brain plots.** Scene vs counting theta activity contrasts from all subjects with MRI with low threshold (20% of max).

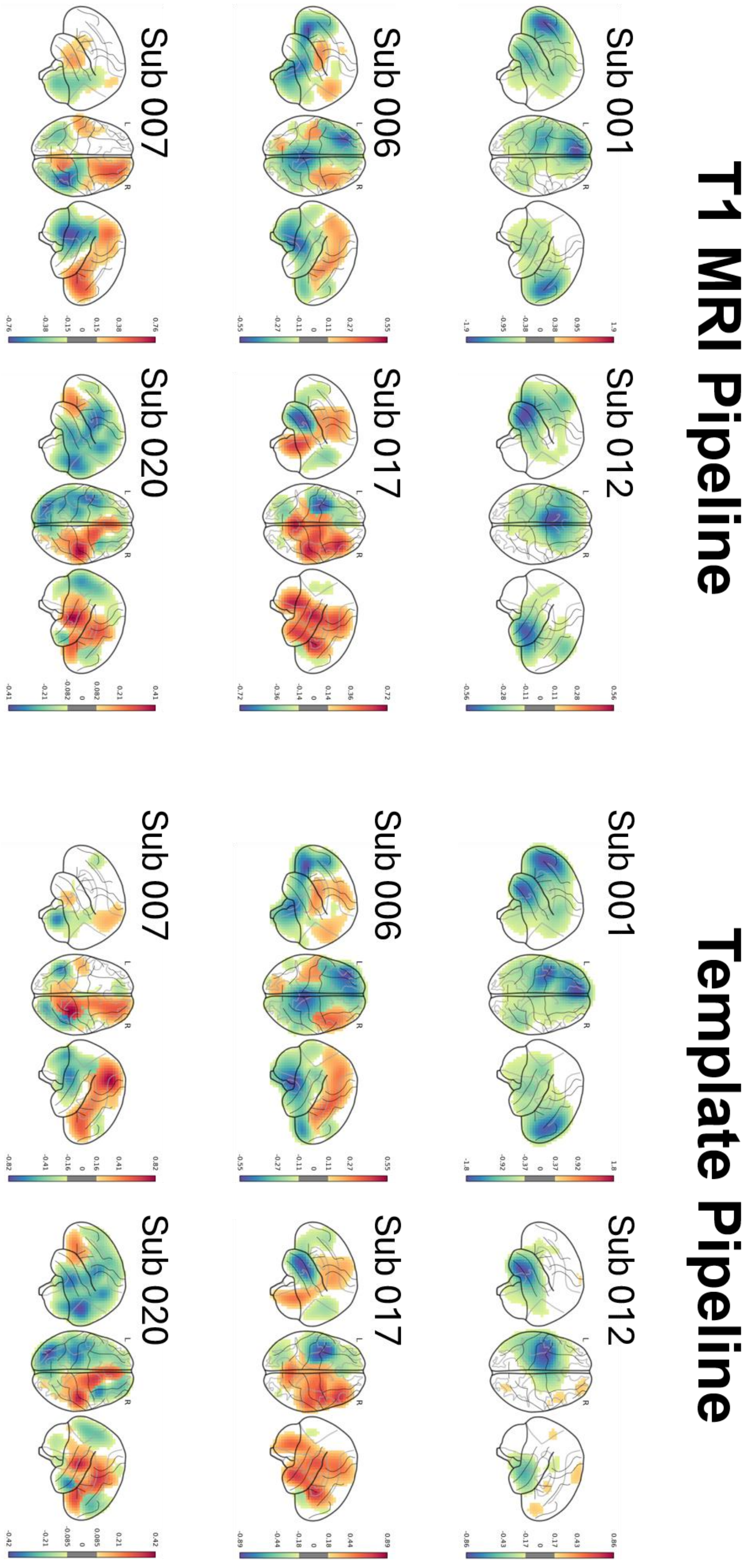

**Supplementary Analysis 1:** To further quantify the correspondence between the two reconstruction models in relation to their activity estimates (dB), we adopted the approach of Rhodes et al. (2025), correlating source-derived estimates between the T1-MRI and template pipelines. We compared peak effect magnitude and peak location across the two pipelines in five regions (left hippocampus, right hippocampus, bilateral hippocampus, temporal-lobe, and whole brain). Across participants (n=6), peak effect magnitude was highly consistent between pipelines in every region (Spearman ρ = 0.943-1; see Table 1), indicating that template-based reconstruction preserved the relative magnitude of the effect across individuals. Peak locations corresponded closely at the median (median separation 6.0-11.7 mm, ~1-2 voxels at 5 mm), though individual subjects showed larger deviations in the broader masks. Peak effect magnitude was strongly correlated between pipelines across participants in all regions (Pearson r = 0.89-1.00, $p < 0.019$; see Supplementary Table 1).

**Supplementary Table 1: Rank Correlation (Spearman) and Pearson's r for peak effects in defined ROIs and whole brain.**

| *Region* | *Spearman Rho* | *Pearson r* | *Median peak distance (mm)* |
|---|---|---|---|
| Left Hippocampus | 0.943 | 0.997 | 6.0 |
| Right Hippocampus | 1 | 0.986 | 9.1 |
| Bilateral Hippocampus | 0.943 | 0.887 | 11.7 |
| Temporal Lobe Mask | 1 | 0.991 | 11.2 |
| Whole Brain | 1 | 0.920 | 9.9 |

**Supplementary Analysis 2:** To assess whether theta-band effects could reflect leakage of alpha power, the source analysis was repeated in the alpha band (8-12 Hz) using an identical pipeline and statistical procedure. No significant clusters were observed in the alpha band (smallest cluster *p* (FWE) = 1.0) across the entire brain (Supplementary Figure 3). We do see some left-lateralised alpha suppression, which could be due to the overlap between the frequency bands at 8Hz. Nevertheless, this in isolation is not significant.

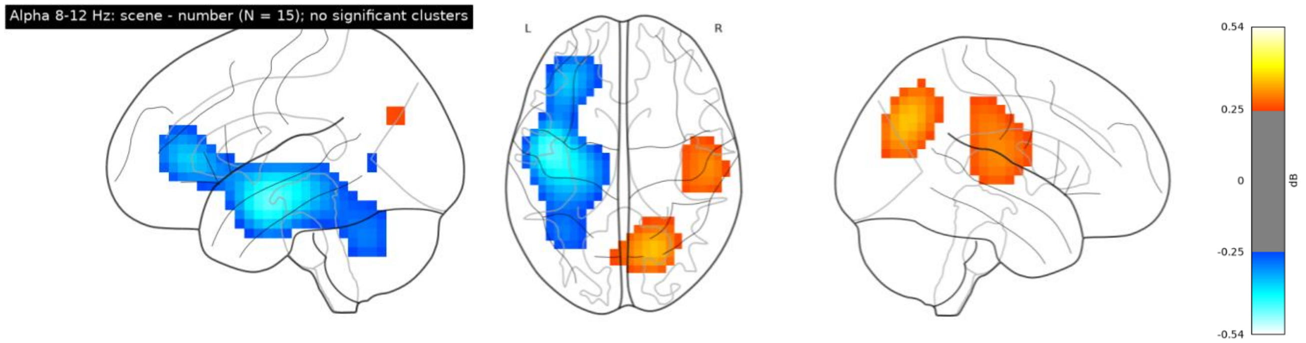


**Supplementary Figure 3:** Group-average scene-versus-number power contrast (dB) in the theta (4-8 Hz, top) and alpha (8-12 Hz, bottom) bands, computed with an identical beamforming pipeline and displayed on a common colour scale. (A) Glass-brain projections. (B) Coronal slices through the medial temporal lobe, with the AAL3 hippocampal boundary outlined in black. No significant alpha-band clusters were observed.
